# Developmental loss of ErbB4 in PV interneurons disrupts state-dependent cortical circuit dynamics

**DOI:** 10.1101/2020.12.09.418590

**Authors:** Renata Batista-Brito, Antara Majumdar, Alejandro Nuno, Martin Vinck, Jessica A. Cardin

## Abstract

GABAergic inhibition plays an important role in the establishment and maintenance of cortical circuits during development. Neuregulin 1 (Nrg1) and its interneuron-specific receptor ErbB4 are key elements of a signaling pathway critical for the maturation and proper synaptic connectivity of interneurons. Using conditional deletions of the *ERBB4* gene in mice, we tested the role of this signaling pathway at two developmental timepoints in parvalbumin-expressing (PV) interneurons, the largest subpopulation of cortical GABAergic cells. Loss of ErbB4 in PV interneurons during embryonic, but not late postnatal, development leads to alterations in the activity of excitatory and inhibitory cortical neurons, along with severe disruption of cortical temporal organization. These impairments emerge by the end of the second postnatal week, prior to the complete maturation of the PV interneurons themselves. Early loss of ErbB4 in PV interneurons also results in profound dysregulation of excitatory pyramidal neuron dendritic architecture and a redistribution of spine density at the apical dendritic tuft. In association with these deficits, excitatory cortical neurons exhibit normal tuning for sensory inputs, but a loss of state-dependent modulation of the gain of sensory responses. Together these data support a key role for early developmental Nrg1/ErbB4 signaling in PV interneurons as powerful mechanism underlying the maturation of both the inhibitory and excitatory components of cortical circuits.

## Introduction

GABAergic interneurons play critical roles in the establishment, maintenance, and mature function of cortical circuits. The diversity of distinct classes of inhibitory interneurons, with different intrinsic properties, morphology, synaptic targeting, and molecular markers, allows them to dynamically sculpt cortical activity during both development and mature function. GABAergic inhibition regulates early organizational activity patterns in the cortex and hippocampus (Allene et al., 2008; Cossart, 2011; Modol et al., 2017) and controls the expression of critical period plasticity by excitatory neurons (Takesian and Hensch, 2013). Developmental dysregulation of GABAergic cells is associated with pathophysiology underlying neurodevelopmental disorders including autism and schizophrenia, as well as epilepsy (Rossignol, 2011). In rodents, the intrinsic properties and synaptic connections of inhibitory interneurons mature over the initial postnatal period, suggesting that early postnatal life may represent a period of GABAergic vulnerability.

The signaling factor Neuregulin-1 (Nrg-1) and its membrane-bound tyrosine kinase receptor ERBB4 (ErbB4) are components of a signaling pathway critical for the proper development of inhibitory neocortical and hippocampal circuits. ErbB4 is selectively expressed by GABAergic interneurons in the neocortex and hippocampus (Yau et al., 2003; Vullhorst et al., 2009; Fazzari et al., 2010; Neddens and Buonanno, 2010; Abe et al., 2011; Del Pino et al., 2013; Batista-Brito et al., 2017). Nrg1/ErbB4 signaling is critical for the formation of inhibitory synapses from parvalbumin-expressing interneurons (PV-INs) onto pyramidal neurons in both hippocampus and neocortex, as well as for formation of excitatory synapses onto PV interneurons (Fazzari et al., 2010; Ting et al., 2011; Del Pino et al., 2013). Loss of ErbB4 from PV-INs leads to altered synaptic input and excitability of these cells (Del Pino et al., 2013). ErbB4 likewise appears to be necessary for the formation of excitatory synapses onto vasoactive intestinal peptide-expressing interneurons (VIP-INs) and their GABAergic synapses onto target cells (Batista-Brito et al., 2017), as well as for synapses to and from hippocampal cholecystokinin-expressing interneurons (CCK-INs) (Del Pino et al., 2017). Nrg1/ErbB4 signaling thus robustly regulates the development of excitatory drive to several populations of GABAergic interneurons and their synaptic input to local targets. However, the precise time period in which this signaling pathway is critical for proper development of inhibitory circuits remains unclear.

In contrast, Nrg1/ErbB4 signaling does not directly affect the maturation of pyramidal neurons or formation of excitatory synaptic inputs to these cells (Chen et al., 2010; Fazzari et al., 2010; Ting et al., 2011; Yin et al., 2013). However, this signaling pathway may nonetheless indirectly regulate the maturation of these cells via effects on GABAergic activity during development. Loss of ErbB4 from PV-INs is reported to cause a loss of dendritic spines in hippocampal pyramidal neurons and some prefrontal cortical neurons (Barros et al., 2009; Del Pino et al., 2013; Yin et al., 2013), suggesting a potential role for Nrg1/ErbB4 control of PV-IN circuits in regulating excitatory synaptic input to pyramidal neurons. However, because most studies have focused on hippocampal neurons, the impact of Nrg1/ErbB4 signaling via PV-INs on cortical pyramidal neurons remains largely unknown.

The disruption of proper synaptic connectivity following loss of Nrg1/EbB4 signaling may impair the function of neocortical and hippocampal circuits. Both global loss and acute inhibition of ErbB4 reduces field potential synchrony between the hippocampus and prefrontal cortex (Tan et al., 2018). Selective loss of ErbB4 from PV-INs likewise leads to broadly disrupted coherence between hippocampus and prefrontal cortex and increased hippocampal gamma oscillations, suggesting disrupted patterns of circuit activity (Del Pino et al., 2013). However, the impact of ErbB4 loss from PV-INs on the firing patterns of local circuits in the intact brain remains unclear.

We generated conditional mutations of *ErbB4* selectively in PV-INs in mice at two key developmental timepoints using the LIM homeodomain transcription factor 6 (Lhx6) promoter to drive prenatal deletion and the parvalbumin (PV) promoter to drive deletion after the second postnatal week. We find that early, but not late, deletion of *ErbB4* causes elevated firing of cortical interneurons and excitatory neurons and perturbation of the temporal pattern of cortical activity that arises early in postnatal life. Early loss of ErbB4 in PV-INs impairs pyramidal neuron dendritic morphology and leads to altered spine density at the apical dendritic tuft. Finally, we find that early deletion of *ErbB4* causes a loss of cortical state-dependent regulation but preserves stimulus feature selectivity for visual inputs.

## Results

### Developmental deletion of ErbB4 from PV interneurons

We directly tested whether developmental dysfunction of PV-INs impairs cortical circuits following disruption of the ErbB4-Nrg1 signaling pathway, using mouse primary visual cortex (V1) as a model for local circuit function. In previous work, we have found that ErbB4 expression in V1 is restricted to GABAergic interneurons and is present in most parvalbumin-(PV) and vasoactive intestinal peptide (VIP)-expressing and a small number of somatostatin-expressing (SST) interneurons (Batista-Brito et al., 2017). To identify whether *ErbB4* disruption in PV interneurons (PV-INs) within an early developmental window has a key impact on cortical maturation, we selectively deleted *ErbB4* from PV-INs at two developmental timepoints by generating *ErbB4^F/F^,Lhx6^Cre^* (Lhx6 mutant) and *ErbB4^F/F^,PV^Cre^* (PV mutant) mice to drive embryonic and late postnatal deletions, respectively. Lhx6 is a transcription factor expressed by neurons originating in the medial ganglionic eminence (MGE), including a majority of PV and SST cortical interneurons (Lavdas et al., 1999; Fogarty et al., 2007). Because ErbB4 is expressed by most PV-INs but few SST interneurons (SST-INs), the Lhx6 mutant mice represent a largely PV-specific embryonic deletion of *ErbB4* (Yau et al., 2003; Del Pino et al., 2013; Batista-Brito et al., 2017). In comparison, because the parvalbumin promoter only becomes active near the end of the second postnatal week (del Rio et al., 1994), the PV mutant mice represent a PV interneuron-specific postnatal deletion of *ErbB4*. To account for any potential impact of the small population of ErbB4-expressing SST-INs, we also generated *ErbB4^F/F^,SST^Cre^* (SST mutant) mice.

### Altered cortical activity in ErbB4^F/F^, Lhx6^Cre^ mutants

To assay the effects of *ErbB4* deletion on cortical activity, we performed extracellular recordings of regular spiking (RS; putative excitatory) neurons and fast spiking (FS; putative PV inhibitory) interneurons throughout cortical layers 2-6 in awake, head-fixed adult mutant and control mice (Figures 1, S1). Early deletion of *ErbB4* from PV-INs in Lhx6 mutants led to marked increases in the spontaneous cortical activity patterns of both RS and FS cells (Figures 1A,B, S1 D). RS cells in Lhx6 mutants exhibited ~5-fold higher spontaneous firing rates than those in controls. Later deletion of *ErbB4* from PV-INs in the PV mutant mice did not affect cortical RS or FS firing rates. Likewise, SST-IN-specific deletion of *ErbB4* in the SST mutant mice did not affect firing rates of RS or FS cells (Figures 1B, S1 D). In parallel with these changes in cortical firing, Lhx6, but not PV or SST, mutants exhibited alterations in local field potential (LFP) activity, with enhanced power in higher frequency bands (>20Hz) during periods of both quiescence and locomotion (Figure 1C, S1 E,F). Together, these data suggest that early, but not late, deletion of *ErbB4* from PV-INs leads to increased firing of cortical excitatory and inhibitory neurons and disrupted cortical activity patterns.

**Figure 1.**
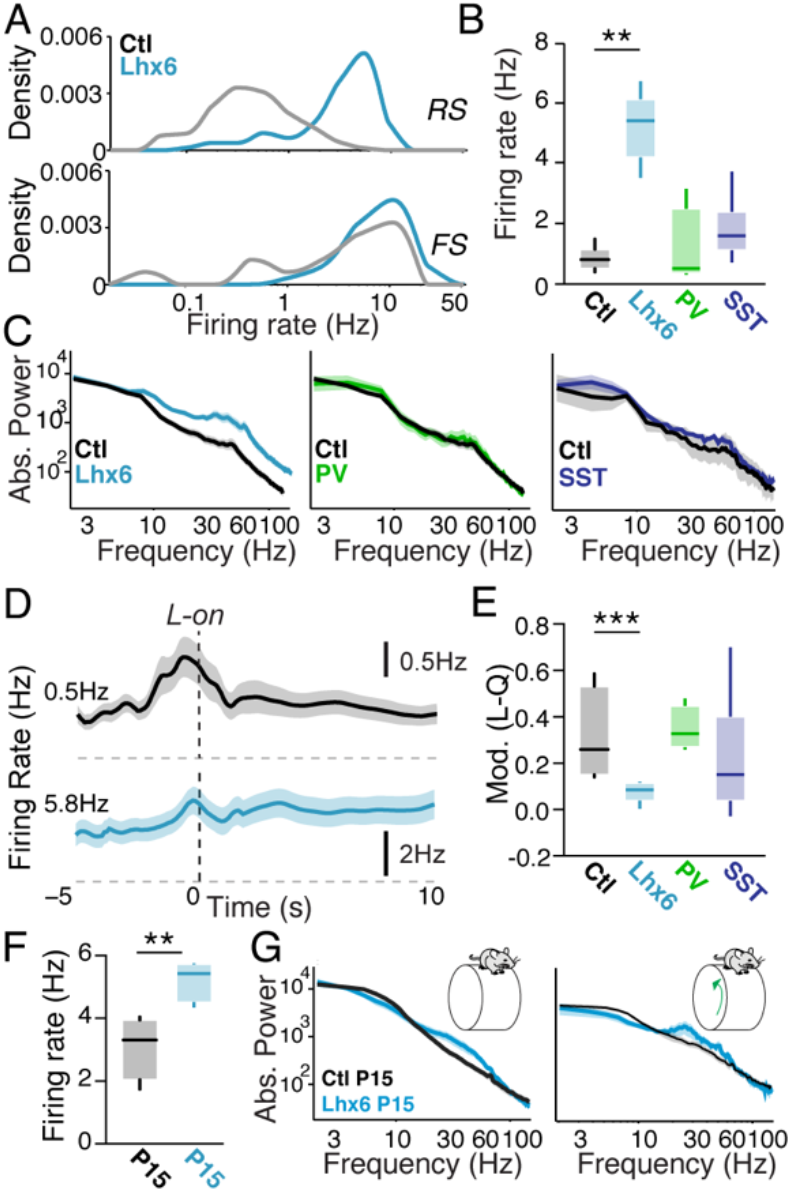
Early, but not late, ErbB4 deletion from PV interneurons alters cortical firing and state transitions. A. Distribution of firing rates across the population of regular spiking (RS, upper panel) and fast-spiking (FS, lower panel) cortical neurons in controls (black) and Lhx6 mutants (cyan). B. Average firing rate of RS cells during quiescence in controls and Lhx6 (cyan), PV (green), and SST (blue) mutants. Controls: 119 cells, 6 mice. Lhx6 mutants: 109 cells, 6 mice. PV mutants: 23 cells, 4 mice. SST mutants: 26 cells, 6 mice. C. LFP power spectra during locomotion for controls and Lhx6, PV, and SST mutants. D. Average firing rate for RS cells around locomotion onset (L-on) in controls (upper panel) and Lhx6 mutants (lower panel). E. Firing rate modulation index (L-Q/L+Q) in early locomotion period (L; −0.5 to 0.5s around L-on) as compared to quiescence period (Q) for RS cells in each group. Controls: 85 cells, 5 mice. Lhx6 mutants: 96 cells, 6 mice. PV mutants: 16 cells, 4 mice. SST mutants: 22 cells, 6 mice. F. Average firing rate for RS cells during quiescence for P15 controls and Lhx6 mutants. Controls: 18 cells, 4 mice. Lhx6 mutants: 19 cells, 6 mice. G. LFP power spectra during quiescence (left) and locomotion (right) for P15 controls and Lhx6 mutants. Shaded areas denote s.e.m.

In previous work, we have identified key GABAergic contributions to state-dependent modulation of cortical activity (Vinck et al., 2015; Batista-Brito et al., 2017; Mossner et al., 2020). We therefore tested the impact of *ErbB4* deletion from PV cells on the ability of cortical circuits to follow transitions in behavioral state. We recorded extracellular signals in V1 cortex of mice transitioning between quiescent and active periods (Vinck et al., 2015; Batista-Brito et al., 2017). In contrast to robust increases in spiking in control animals, RS cells in the Lhx6 mutants showed reduced changes in firing at locomotion onset (Figure 1D,E). Likewise, RS cells in the Lhx6 mutants showed little change at locomotion offset, a separate period of high global arousal that is independent of motor activity and normally associated with decreased firing rates (Figure S1G) (Vinck et al., 2015). FS cells in the Lhx6 mutants did not exhibit impaired state modulation (Figure S1H, I). Loss of ErbB4 in the PV and SST mutants had no impact on state-dependent modulation of cortical firing rates of RS (Fig 1E) or FS (Fig S1I) cells. These data indicate a reduction of the overall cortical response to behavioral arousal in the Lhx6 mutants and suggest that early, but not late, *ErbB4* deletion from PV-INs and subsequent disruption of GABAergic synaptic transmission decreases the impact of locomotion- and arousal-related signals on cortical excitatory neurons.

To more directly assay the developmental impact of early *ErbB4* deletion from PV-INs, we performed extracellular recordings of cortical activity in awake, head-fixed controls and Lhx6 mutants on postnatal day 15. Early loss of ErbB4 in the Lhx6 mutants resulted in increased firing rates of RS cells in the young mutants as compared to controls (Figure 1F). In parallel, young Lhx6 mutants exhibited enhanced LFP activity in higher frequency bands (Figure 1G). Together, these data suggest that the loss of ErbB4 from PV-INs exerts a detrimental impact on cortical function that emerges prior to the full maturation of PV-IN firing properties (Goldberg et al., 2011).

### Perturbation of key forms of temporally patterned cortical activity in ErbB4^F/F^,Lhx6^Cre^ mutants

Neuronal synchronization is a core feature of cortical activity and altered neural rhythms are a hallmark neurodevelopmental disorders, including schizophrenia (Uhlhaas and Singer, 2010; Gandal et al., 2012). To examine the impact of *ErbB4* deletion in PV interneurons on the temporal patterning of cortical activity, we assessed the relationship between spiking activity and LFPs in V1 (Figure 2). The Lhx6 mutants showed a reduction in the phase-locking of individual RS cells to low frequencies (1-6Hz) and gamma-range (40-60Hz) LFP oscillations (Figure 2B,C). In contrast, FS cells in the Lhx6 mutants exhibited a loss of phase-locking to low frequencies and a robust increase in phase-locking to gamma frequencies during locomotion (Figure 2B,C) and quiescence (Figure S2). Correlations between the spiking of simultaneously recorded RS-RS pairs were substantially diminished in mutants (Figure 2E), indicating a loss of synchronous firing. RS-FS synchrony was likewise decreased in mutants (Figure S2). FS cells typically exhibit a high level of synchrony with other FS cells, largely due to dense chemical and electrical synaptic connectivity (Gibson et al., 1999). FS-FS synchrony was unaffected in mutants (Figure 2E). These data indicate that early dysregulation of PV-INs reduces the entrainment of excitatory activity and impairs excitatory synchrony in cortical networks, disrupting several key forms of temporally organized activity that are important for information processing in cortical circuits (Fries, 2009).

**Figure 2.**
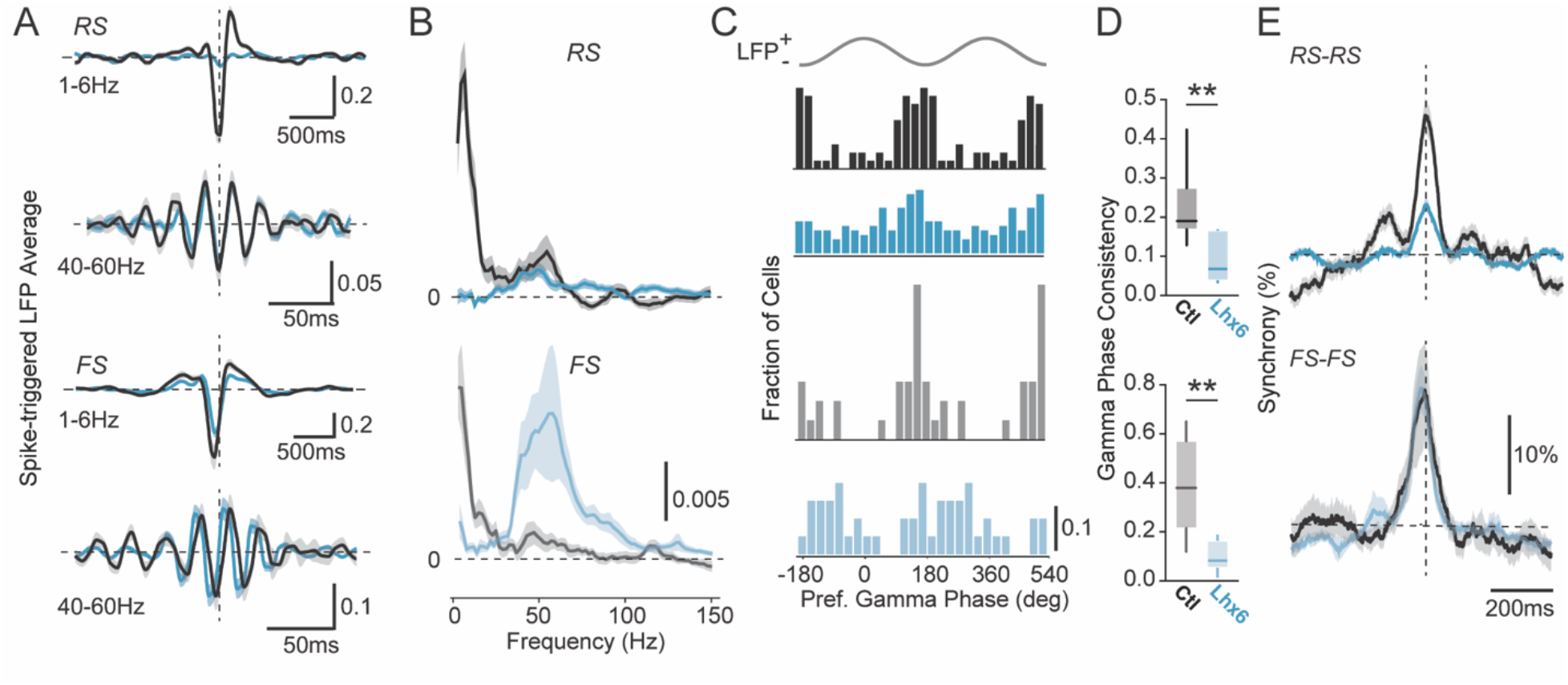
Loss of PV ErbB4 disrupts the temporal organization of cortical activity. A. Spike-triggered LFP average in 1-6 Hz and 40-60Hz bands during locomotion for controls (black) and Lhx6 mutants (cyan). B. Average spike-LFP phase-locking during locomotion. C. Preferred LFP gamma-phase of firing during locomotion for RS (upper) and FS (lower) cells. D. Consistency of preferred LFP gamma-phases for RS (upper) and FS (lower) cells. Controls: 130 RS cells, 16 FS cells, 6 mice. Lhx6 mutants: 117 RS cells, 31 FS cells, 6 mice. E. Average normalized cross-correlograms during quiescence for RS-RS (upper) and FS-FS (lower) pairs. Controls: 192 RS-RS pairs, 15 FS-FS pairs, 6 mice. Shaded areas denote s.e.m.

### Early loss of ErbB4 in PV interneurons impairs pyramidal neuron morphology

GABAergic activity patterns in developing cortical circuits play a powerful role in regulating the maturation of synaptic connectivity and neuronal function (Fazzari et al., 2010; Fishell and Rudy, 2011; Tuncdemir et al., 2016; Batista-Brito et al., 2017; Takesian et al., 2018). To identify cellular changes that may underlie the circuit-level disruption we observed in the mutants, we examined the impact of early developmental loss of ErbB4 in PV-INs on the maturation of pyramidal cell dendritic morphology. We reconstructed pyramidal neurons in cortical layers 2/3 and 5 of both mutants and controls (Figure 3, S3). Lhx6 mutants showed substantial reductions in the dendritic length and branching of layer 5 pyramidal neurons (Figure 3B-E), and these effects were observed in both apical and basal dendrites (Figure S3 H-J). Layer 2/3 pyramidal cells in the Lhx6 mutants exhibited similar decreases in dendritic length and branching (Figure S3).

**Figure 3.**
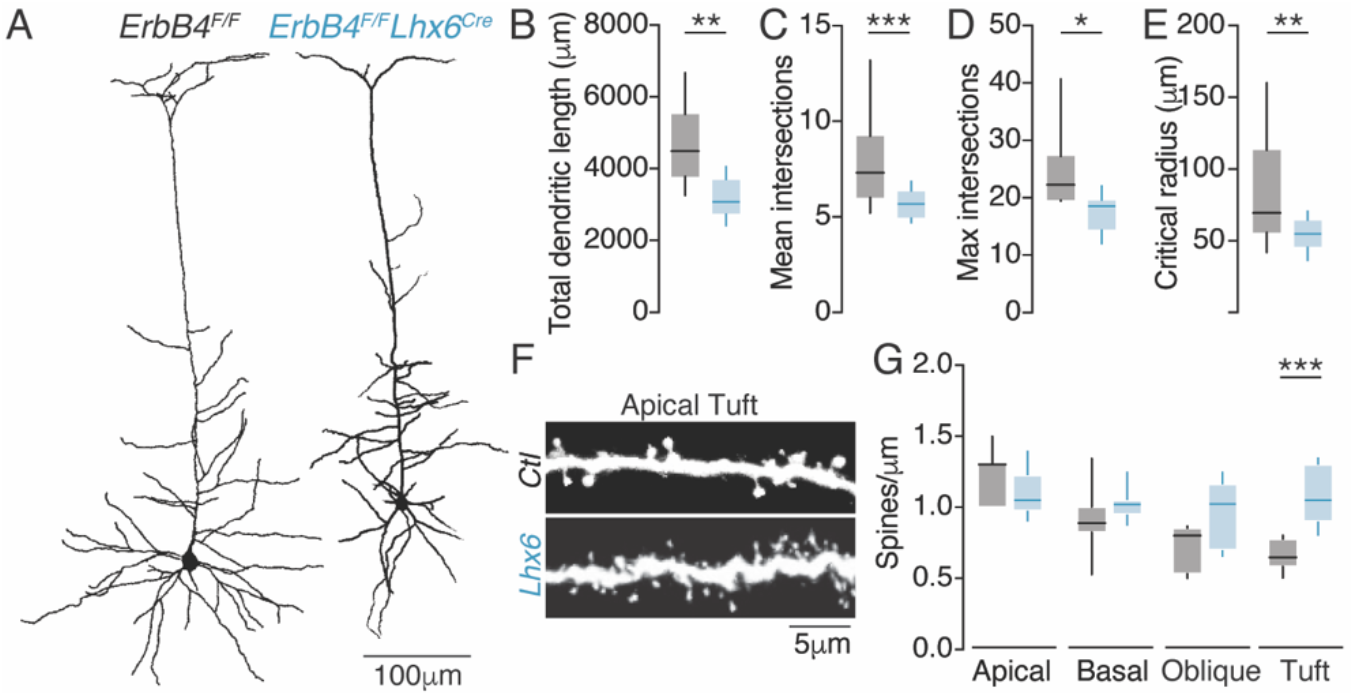
PV ErbB4 deletion impairs excitatory pyramidal neuron morphology. A. Example layer 5 pyramidal neuron reconstructions from an *ErbB4^F/F^* control (left) and an *ErbB4^F/F^,Lhx6^Cre^* mutant (right). B. Average total dendritic length for layer 5 pyramidal neurons in controls (black) and Lhx6 mutants (cyan). Controls: 15 cells. Lhx6 mutants: 13 cells. C. Average mean dendritic intersections for control and Lhx6 mutant layer 5 cells. D. Average maximum dendritic intersections for control and Lhx6 mutant layer 5 cells. E. Average critical radius for control and Lhx6 mutant layer 5 cells. F. Example spine density at the apical dendritic tuft of layer 5 pyramidal neurons in controls and Lhx6 mutants. G. Spine density on the main apical dendrite, basal dendrites, oblique dendrites, and apical dendritic tuft for layer 5 pyramidal neurons in controls and Lhx6 mutants. Controls: 10 cells. Lhx6 mutants: 10 cells.

Because dendrites are the major site of excitatory synaptic input to pyramidal neurons, we further examined the impact of disrupted Nrg1/ErbB4 signaling on excitatory synaptic inputs by measuring the distribution of dendritic spines. Layer 5 pyramidal neurons in the Lhx6 mutants exhibited an altered dendritic spine distribution, with a significant increase in spine density in the apical dendritic tuft (Figure 3 F,G, S3 F). Together, these data suggest that early loss of ErbB4 in PV interneurons reduces excitatory neuron dendritic complexity across cortical layers and leads to abnormally high spine density at the apical tuft of layer 5 pyramidal neurons.

### Prenatal disruption of PV interneurons impairs state-dependent modulation of sensory processing

Previous work has shown that the visual tuning of mouse V1 pyramidal neurons arises from feedforward thalamic inputs and is amplified by cortico-cortical inputs (Lien and Scanziani, 2013). However, visual response gain is also modulated by feedforward and feedback inputs that target apical dendrites and provide multimodal contextual information (Jia et al., 2010; Cruz-Martin et al., 2014; Roth et al., 2016; Ranganathan et al., 2018), along with inhibitory dendritic inputs that are modulated by arousal and locomotion (Fu et al., 2014a; Zhang et al., 2014; Pakan et al., 2016; Dipoppa et al., 2018). To determine the impact of ErbB4 loss on the tuning and state-dependent gain components of pyramidal neuron visual responses, we therefore performed extracellular recordings in V1 of awake behaving animals during visual stimulation (Figures 4, S4). In the Lhx6 mutants, RS cell visual responses to drifting grating stimuli were reduced in amplitude (Figure 4A,B) but exhibited normal orientation selectivity (Figure 4C). FS visual responses were likewise decreased in amplitude (Figure S4C, D). Neurons in controls and Lhx6 mutants both showed bias towards horizontal (0° angle) stimuli, a feature of cortical visual tuning that is refined by experience-dependent plasticity after eye opening (Rochefort et al., 2011) (Figure 4D, S4 A,B). However, deletion of *ErbB4* from PV-INs eliminated the increase in visual response gain normally observed during periods of locomotion and arousal. We and others have previously shown that the signal-to-noise of visual responses in controls increases in the active state (Niell and Stryker, 2010; Fu et al., 2014b; Vinck et al., 2015), but this enhancement was absent in the mutants (Figure 4E).

**Figure 4.**
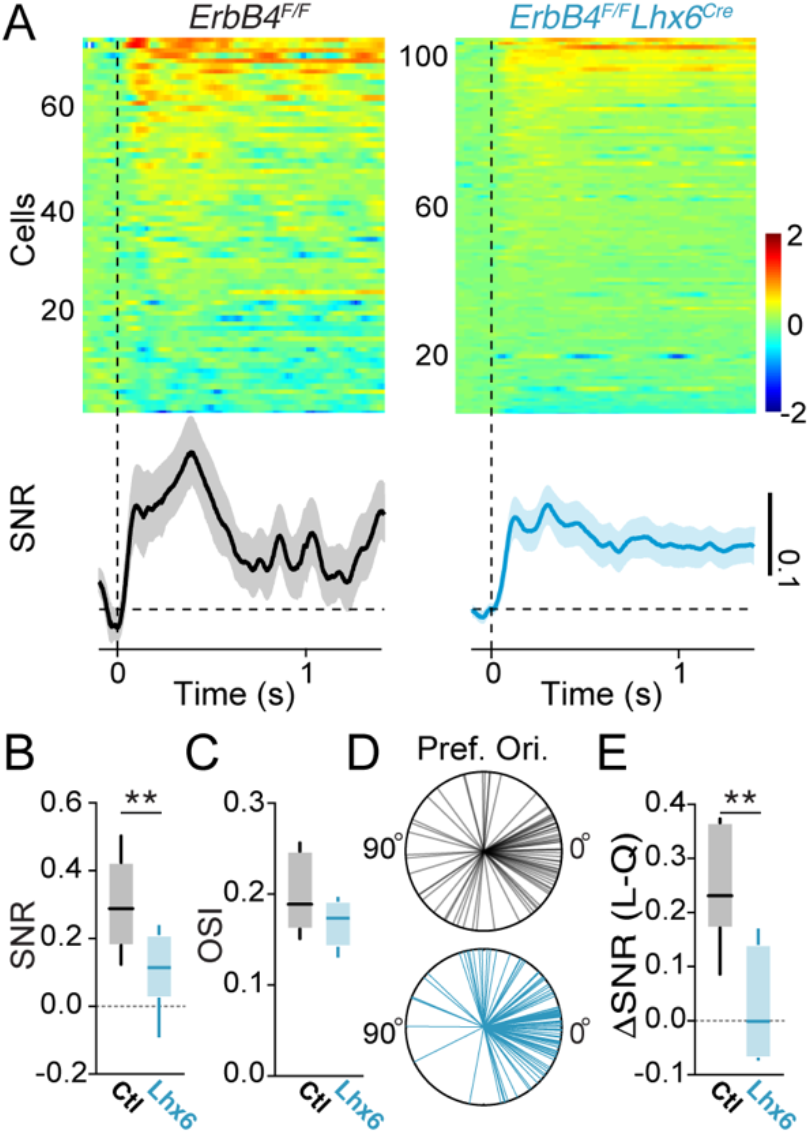
ErbB4 mutants exhibit reduced visual response modulation but intact selectivity. A. Upper panel: visual responses of individual RS cells in controls (left) and Lhx6 mutants (right). Lower panel: average visual response in each population, shown as signal:noise ratio (SNR) over the stimulus presentation period. B. Average signal-to-noise ratio of visual responses for RS cells in controls (black) and Lhx6 mutants (cyan). Controls: 92 cells, 6 mice. Lhx6 mutants: 140 cells, 8 mice. C. Average orientation selectivity index (OSI) of RS cells in controls and Lhx6 mutants. Controls: 83 cells, 6 mice. Lhx6 mutants: 108 cells, 6 mice. D. Radial plots of preferred orientations of all RS cells in each group. E. Increase in stimulus rate modulation of RS cells during locomotion as compared to quiescence. Controls: 86 cells, 6 mice. Lhx6 mutants: 100 cells, 6 mice.

## Discussion

Our results reveal a key role for Nrg1/ErbB4 signaling in PV-INs early in neocortical development. We examined the consequences of conditional *ErbB4* deletion from PV-INs at two developmental periods, using promoters that become active at prenatal and late postnatal timepoints. We find that early *ErbB4* deletion from PV-INs disrupts the firing patterns of both putative fast-spiking interneurons and excitatory neurons and leads to impaired state-dependent function of neocortical circuits. We also find that loss of ErbB4 from PV-INs impairs the dendritic morphology of neocortical pyramidal neurons and alters spine density, suggesting that the circuit-level deficits observed following selective developmental perturbation of GABAergic inhibition arise from impairments of the synaptic connectivity and firing patterns of both inhibitory and excitatory neurons.

Recordings in primary visual cortex of awake mice revealed a robust impact of early *ErbB4* deletion on the pattern of cortical activity. We found that early loss of ErbB4 from PV-INs in the Lhx6 mutants, but not late loss of ErbB4 in the PV mutants, caused a ~5-fold increase in the firing rates of putative excitatory cells in the adult cortex. Early loss of ErbB4 likewise led to increased firing rates of fast-spiking, putative PV-INs. These data are in good agreement with *ex vivo* data from the hippocampus, where fast-spiking interneuron firing rates are elevated following deletion of *ErbB4* (Del Pino et al., 2013). The Lhx6 mutants, but not the PV mutants, also exhibited altered cortical gamma-range LFP activity, which is associated with cognition and sensory processing (Cardin, 2016, 2018) and requires the proper synaptic interaction between PV-INs and excitatory neurons (Cardin et al., 2009). Enhanced gamma-range activity has also previously been observed in the hippocampus in *ErbB4^F/F^,Lhx6^Cre^* mutants (Del Pino et al., 2013). Deletion of *ErbB4* from somatostatin-expressing interneurons had no effect on cortical firing rates or LFP activity.

Cortical activity patterns and sensory responses are strongly modulated by behavioral state, such as sleep, wakefulness, and attention. This modulation is thought to be important for enhancing encoding of behaviorally relevant information and is compromised in disease (Lewis and Lieberman, 2000). In the healthy cortex, transitions between behavioral states, such as quiescence and arousal or locomotion, are associated with robust modulation of cortical firing rates (Niell and Stryker, 2010; McGinley et al., 2015; Vinck et al., 2015; Tang and Higley, 2020). Previous work has suggested that dendrite-targeting interneurons, the VIP- and SST-expressing cells, play key roles in mediating state-dependent modulation of cortical circuit activity (Fu et al., 2014b; Pakan et al., 2016; Dipoppa et al., 2018; Urban-Ciecko et al., 2018). Surprisingly, early loss of ErbB4 in the Lhx6 mutants, but not late loss in the PV mutants, caused a dysregulation of this state-dependent modulation. These results suggest an unanticipated role for PV-INs in cortical state modulation, potentially via synaptic interactions with other GABAergic populations (David et al., 2007; Pfeffer et al., 2013; Pi et al., 2013; Walker et al., 2016; Cardin, 2018).

To more closely examine the onset of cortical disruption following loss of ErbB4, we recorded cortical activity in awake mice at the end of the second postnatal week. We found that at P15 the Lhx6 mutants exhibited elevated cortical firing rates and altered gamma-range LFP activity similar to that observed in adults, suggesting that loss of Nrg1/ErbB4 signaling in PV-INs causes key impairments in cortical circuit function that emerge by the end of the second postnatal week. These effects arise before the maturation of the intrinsic and fast-spiking properties of the PV-INs, which occurs rapidly after this timepoint (Goldberg et al., 2011).

Tightly coupled, fine time-scale interactions between excitation and inhibition are critical for restricting the timing and number of excitatory action potentials (Pouille and Scanziani, 2001; Wehr and Zador, 2003; Cardin et al., 2010) and for information encoding (Fries, 2009). Synchrony between cells may enhance the transmission of information across long-range cortical circuits. In healthy cortical circuits, excitatory neurons are entrained to the gamma rhythm and their activity is tightly coupled to that of fast-spiking inhibitory interneurons (Hasenstaub et al., 2005; Cardin et al., 2009; Vinck et al., 2013; Cardin, 2016). Following deletion of *ErbB4* from PV-INs in the Lhx6 mutants, RS cells showed a loss of phase consistency in their entrainment to the gamma rhythm and exhibited a near-complete loss of pairwise synchrony. RS cells in the mutants also exhibited diminished synchrony with FS cells. In contrast, FS cells showed an abnormal enhancement of entrainment to the gamma rhythm and no impairment of pairwise synchrony. Early loss of Nrg1/ErbB4 signaling in PV-INs thus dysregulates the fine temporal structure of cortical activity and decouples the firing of RS and FS cells, degrading the ability of cortical circuits to participate in both the encoding and transmission of information. Loss of ErbB4 in VIP-INs, the other major group of ErbB4-expressing neocortical interneurons, likewise causes a loss of RS cell coupling to the gamma rhythm and decreased synchrony (Batista-Brito et al., 2017), suggesting that both populations of GABAergic interneurons are critical to the maturation of cortical circuit organization and the temporal patterning of cortical activity.

Previous work has suggested that disruption of Nrg1/ErbB4 signaling may affect the formation of excitatory inputs to pyramidal neurons. Nrg1 application does not affect excitatory synaptic development on cortical or hippocampal pyramidal neurons (Ting et al., 2011), and disruption of Nrg1 signaling does not affect pyramidal neuron migration (Fazzari et al., 2010). Deletion of *ErbB4* from pyramidal neurons by genetic or viral means does not affect synaptic development or morphology of these cells (Chen et al., 2010; Fazzari et al., 2010; Ting et al., 2011; Yin et al., 2013). Likewise, neither genetic deletion nor overexpression of *ErbB4* in pyramidal neurons alters spine density or excitatory synapse function in hippocampal pyramidal neurons (Yin et al., 2013). In contrast, late deletion of *ErbB4* from PV-INs leads to reduced spine density and decreased miniature excitatory post-synaptic current (EPSC) frequency in hippocampal pyramidal neurons, consistent with a decrease in excitatory synaptic input (Yin et al., 2013). However, the effects of this mutation on cortical pyramidal neurons are less clear, as spine density is reduced only in a subset of frontal cortical neurons (Yin et al., 2013). Early conditional deletion of *ErbB4* in Lhx6 mutants likewise causes a loss of spines in hippocampal pyramidal neurons, but an increase in spontaneous EPSCs (Del Pino et al., 2013). However, the effects of early *ErbB4* deletion on neocortical neurons remain unclear.

To examine the impact of disrupted Nrg1/ErbB4 signaling on neocortical excitatory neurons, we assessed the morphology of pyramidal neurons in superficial and deep cortical layers of primary visual cortex in the Lhx6 mutants. We found that early perturbation of Nrg1/ErbB4 signaling in PV-INs led to a substantial loss of dendritic length and complexity in excitatory pyramidal neurons in layers 2/3 and 5. In layer 5 pyramidal neurons, we further observed a redistribution of dendritic spines, with increased spine density at the apical dendritic tuft. These data suggest that early disruption of PV-IN synaptic transmission following loss of ErbB4 causes impairments in the maturation or maintenance of pyramidal neuron dendrites and excitatory synaptic connections at spines. Our findings thus highlight a robust regulation of the morphology and apical dendritic excitatory synaptic connectivity of cortical excitatory neurons via Nrg1/ErbB4 signaling in GABAergic interneurons.

Excitatory neurons in V1 exhibit robust tuning for visual features that arises from thalamocortical inputs and is amplified by local cortico-cortical connections between cells with similar tuning (Ko et al., 2011; Lien and Scanziani, 2013). In addition, the apical dendritic tufts of cortical pyramidal neurons in layer 1 receive top-down feedback inputs from higher-order cortical areas (Rockland and Pandya, 1979; Felleman and Van Essen, 1991; Cauller et al., 1998) and inputs from thalamocortical projections (Roth et al., 2016). These long-range excitatory inputs are thought to provide contextual information to primary cortex neurons, regulating the integration and gain of sensory-evoked responses (Schiller et al., 1997; Larkum and Zhu, 2002; Larkum et al., 2004; Wang et al., 2007; Nassi et al., 2013; Keller et al., 2020). In addition, pyramidal neurons also receive inhibitory inputs on both apical dendritic shafts and spines (refs) that are robustly modulated by arousal and locomotion (Fu et al., 2014a; Pakan et al., 2016; Dipoppa et al., 2018) and can modulate excitatory synaptic inputs to spines (Chiu et al., 2013). We tested the impact of ErbB4 deletion on the tuning and gain regulation of visual responses of V1 pyramidal neurons across behavioral states. Following deletion of *ErbB4* from PV-INs, RS cells in visual cortex exhibited normal feature selectivity for stimulus orientation, suggesting largely intact feedforward thalamocortical input. In the normally developing cortex, the gain of visual responses is strongly modulated by behavioral state and enhanced during locomotion (Niell and Stryker, 2010; Bennett et al., 2013; Vinck et al., 2015). However, state-dependent gain modulation of visual responses was eliminated in the mutants, suggesting perturbation of the feedforward and feedback inputs to apical dendrites that provide state-dependent and contextual input.

In summary, we find that embryonic, but not late postnatal, Nrg1/ErbB4 signaling is critical for the proper maturation of cortical circuits. Early loss of ErbB4 in PV interneurons leads to profound dysregulation of firing and a loss of temporal patterning of cortical circuit activity as early as the second postnatal week. In parallel, disruption of Nrg1/ErbB4 signaling in PV interneurons also results in impaired pyramidal neuron morphology and apical dendritic spine distribution, suggesting indirect regulation of pyramidal cell maturation or maintenance via control of GABAergic signaling. In association, early deletion of *ErbB4* in PV interneurons does not affect visual response tuning, but results in loss of state-dependent modulation of visual response gain. Early developmental Nrg1/ErbB4 signaling in PV interneurons is thus a powerful mechanism underlying the establishment of proper excitatory-inhibitory circuit interactions and state-dependent sensory processing in the cortex.

## Author Contributions

RBB, MV, AM, AN, and JAC designed experiments. RBB, MV, and JAC performed *in vivo* electrophysiology recordings. RBB, MV, and JAC analyzed *in vivo* electrophysiology data. AM and AN performed virus injections and cellular reconstructions and analyzed anatomical data. RBB, AM, AN, and JAC wrote the manuscript.

## Acknowledgements

The authors thank Dr. M.J. Higley for helpful comments on the manuscript and Dr. A. Koleske for the ErbB4^F/F^ mice. This work was supported by a Brown-Coxe fellowship, a Jane Coffin Childs Fellowship, a NARSAD Young Investigator Award, and a Simons Bridge to Independence Award to RBB; a Rubicon fellowship and a Human Frontiers Postdoctoral Fellowship to MV; and NIH R01 MH102365, NIH R01 EY022951, a Smith Family Award for Excellence in Biomedical Research, a Klingenstein Fellowship Award, an Alfred P. Sloan Fellowship, a NARSAD Young Investigator Award, and a McKnight Fellowship to JAC. This work was further supported by the Yale Core Grant for Vision Research NIH P30 EY02678.

## Methods

All animal handling and maintenance was performed according to the regulations of the Institutional Animal Care and Use Committee of the Yale University School of Medicine. *ErbB4^F/F^* mice were crossed to *ErbB4^F/+^*,*Lhx6^Cre^, ErbB4^F/+^*,*PV^Cre^*, or *ErbB4^F/+^*,*SST^Cre^* to generate mutant animals (*ErbB4^F/F^*,*Lhx6^Cre^; ErbB4^F/F^*,*PV^Cre^; ErbB4^F/F^*,*SST^Cre^*) and littermate controls (*ErbB4^F/F^*). In a subset of reconstruction experiments, wild-type c57Bl/6 animals were used. We used both female and male animals ranging from ages P15 to P90, as specified in the experimental descriptions.

### Immunohistochemistry and anatomical reconstructions

Layer 2/3 and Layer 5 pyramidal neurons were labeled in control mice using a Cre-dependent viral labeling method. AAV-CamKII-Cre virus (Addgene) was diluted 1: 10^6 in 1X cold phosphate buffer solution. This was mixed with an AAV-CAG-Flex-tdTomato (Addgene) virus in a 50:50 mix and injected into the visual cortex. The injection was made at a depth of 300 to 500 μm to ensure proper labelling of Layer 2/3 and Layer 5 pyramidal neurons. The virus was allowed to express for 10 days before tissue collection. Layer 2/3 and Layer 5 neurons were labelled in the mutants using a Flp-dependent viral approach. AAV-ef1a-FlpO virus was diluted 1:1,000 in 1X cold phosphate buffer solution. This was mixed with an AAV-ef1a-DIO-eYFP virus in a 50:50 mix and injected in the cortex. The virus was allowed to express for up to three weeks before collection of tissue for analysis.

Following expression of cell label, animals were perfused transcardially with 4% paraformaldehyde (PFA) in 1X phosphate buffered saline (PBS). This was followed by overnight tissue post-fixation in 4% PFA in 1X PBS stored in 4⁰ C. Brains were rinsed in 1X PBS and vibratome sections were obtained at 150 to 200 μm thickness. Nuclear counterstaining was performed with DAPI solution.

Images of neurons were acquired using Zeiss 880 confocal imaging software (Zen Black) at 20x magnification. Stitched Z-stacks of the labelled cells were then imported into Fiji (ImageJ) and traced using the semi-automated plugin Simple Neurite Tracer. We obtained a three-dimensional wire diagram for each neuron, along with a total dendritic length measurement and a ratio of total apical to total basal dendrite length. Simple Neurite Tracer was used to perform a Sholl analysis of each neuron.

For spine counts, Z-stack images of the imaged neurons were collected at 63x and processed in Fiji. Dendritic segments fitting four categories were imaged for each neuron: stretches of primary apical dendrites, basal dendrites (no more than 40 μm from the soma), oblique dendrites off the apical dendrite (more than 40 μm from the soma), and apical dendritic tuft in cortical layer 1. Once the dendrite of interest was localized and categorized, a 10 μm segment was measured and spines were counted.

### Headpost surgery and wheel training

Mice were handled for −10 min/day for 5 days prior to the headpost surgery. On the day of the surgery, the mouse was anesthetized with isoflurane and the scalp was shaved and cleaned three times with Betadine solution. An incision was made at the midline and the scalp resected to each side to leave an open area of skull. Two skull screws (McMaster-Carr) were placed at the anterior and posterior poles. Two nuts (McMaster-Carr) were glued in place over the bregma point with cyanoacrylate and secured with C&B-Metabond (Butler Schein). The Metabond was extended along the sides and back of the skull to cover each screw, leaving a bilateral window of skull uncovered over primary visual cortex. The exposed skull was covered by a layer of cyanoacrylate. The skin was then glued to the edge of the Metabond with cyanoacrylate. Analgesics were given immediately after the surgery and on the two following days to aid recovery. Mice were given a course of antibiotics (Sulfatrim, Butler Schein) to prevent infection and were allowed to recover for 3-5 days following implant surgery before beginning wheel training.

Once recovered from the surgery, mice were trained with a headpost on the wheel apparatus. The mouse wheel apparatus was 3D-printed (Shapeways Inc.) in plastic with a 15 cm diameter and integrated axle and was spring-mounted on a fixed base. A programmable magnetic angle sensor (Digikey) was attached for continuous monitoring of wheel motion. Headposts were custom-designed to mimic the natural head angle of the running mouse, and mice were mounted with the center of the body at the apex of the wheel. On each training day, a headpost was attached to the implanted nuts with two screws (McMaster-Carr). The headpost was then secured with thumb screws at two points on the wheel. Mice were headposted in place for increasing intervals on each successive day. If signs of anxiety or distress were noted, the mouse was removed from the headpost and the training interval was not lengthened on the next day. Mice were trained on the wheel for up to 7 days or until they exhibited robust bouts of running activity during each session. Mice that continued to exhibit signs of distress were not used for awake electrophysiology sessions.

### *In vivo* electrophysiology

All extracellular single-unit, multi-unit, and LFP recordings were made with an array of independently moveable tetrodes mounted in an Eckhorn Microdrive (Thomas Recording). Signals were digitized and recorded by a Digital Lynx system (Neuralynx). All data were sampled at 40kHz. All LFP recordings were referenced to the surface of the cortex. LFP data were recorded with open filters and single unit data was filtered from 600-9000Hz. Awake recordings were made from mice that had received handling and wheel training as described above. On the initial recording day, a small craniotomy was made over V1 under light isoflurane anesthesia. The craniotomy was then covered with Kwik-Cast (World Precision Instruments), after which the mouse was allowed to recover for 2 hours. Mice were then fitted with a headpost and secured in place on the wheel apparatus before electrodes were lowered. Recording electrodes were initially lowered to ~150 μm, then independently adjusted after a recovery period of 30-60 min. At the end of a recording session, the craniotomy was flushed with saline and capped. On subsequent recording days, the craniotomy was flushed with saline before placing the electrode array in a new site. Recordings were performed mainly in the second half of the light portion of the light/dark cycle.

### Visual stimulation

Visual stimuli were presented on an LCD monitor at a spatial resolution of 1680×1050, a real-time frame rate of 60Hz, and a mean luminance of 45 cd/m^2^ positioned 15cm from the eye. The LCD monitor used for visual stimulation (22 inches) was mounted on an arm and positioned on the right side of the animal, perpendicular to the surface of the right eye. The screen was placed so that stimuli were only presented to the right eye. Initial hand-mapping was performed to localize the receptive fields of identified cells in the electrophysiological experiments; an automated mapping was using during imaging to identify the largest overall change in fluorescence for a given field. To maximize data collection, visual stimuli were positioned to cover as many identified receptive fields as possible. All stimuli were sinusoidal drifting gratings at a temporal frequency of 2 Hz, presented at a fixed duration of 1.5 s (2 s) with an interstimulus interval of 2 s (5 s) for electrophysiological (imaging) experiments. For the electrophysiological recordings, we used blocks of visual stimuli where contrast was held at 100% and orientation was varied. To determine orientation tuning, gratings were presented at 12 different orientations, randomized and presented 20-50 times per orientation. Orientation tuned stimuli were optimized for mean spatial frequency.

## Quantification and Statistical Analysis

### In vivo electrophysiology analysis

Spikes were clustered semi-automatically using the following procedure. We first used the KlustaKwik 2.0 software to identify a maximum of 30 clusters using the waveform Energy and Energy of the waveform’s first derivative as clustering features. We then used a modified version of the M-Clust environment to manually separate units and we selected well-isolated units. We further ensured that maximum contamination of the ISI (Inter-spike-interval) histogram <1.5 ms was smaller than 0.1%. In a small number of cases we accepted clusters with isolation distances smaller than 20, which could be caused by e.g. non-Gaussian clusters, only if separation was of sufficient quality as judged by comparing the cluster with all other noise clusters (20%, 80% quantiles of ID = 19, 43; median±standard error of median = 25±0.5). Unit data were analyzed for firing rates and patterns, correlations, and visual responses using custom-written Matlab software.

### Quantification of firing rate

The firing rate was computed by dividing the total number of spikes a cell fired in a given period by the total duration of that period.

### Computation of wheel position and change points

Wheel position was extracted from the output of a linear angle detector. Since wheel position is a circular variable, we first transformed the sensor data to the [−π,π] interval. Because the position data would make sudden jumps around values of π and –π, we further performed circular unwrapping of the position phases to create a linear variable.

We then used a change-point detection algorithm that detected statistical differences in the distribution of locomotion velocities across time. The motivation of this method relative to the standard method of using an arbitrary threshold (e.g., 1 cm/s) (Niell and Stryker, 2010) is that our technique allowed for small perturbations in locomotion speed to be identified that might otherwise fail to reach the locomotion threshold. Further, it ensured that the onset of locomotion could be detected before the speed reached 1 cm/s. If the distributions of data points 100 ms before and 100 ms after a certain time point *t* were significantly different from each other, using a standard T-test at p<0.05 and sampling at 2 kHz, then the data point was deemed a candidate change point. A point *t* was considered a candidate locomotion onset point if the speed 100 ms after *t* was significantly higher than 100 ms before *t*. A point was considered a candidate locomotion offset point if the speed 100 ms after *t* was significantly lower than 100 ms before *t*. A point was accepted as a locomotion onset point if the previous transition point was a locomotion offset point. A point was considered to be a locomotion offset if it was preceded by a locomotion onset point, and if the speed 100 ms after *t* did not significantly differ from zero. This prevented a decrease in speed to be identified as a locomotion offset point. We further required that a locomotion offset point not be followed by a locomotion onset point for at least 2 s, because mice sometimes showed brief interruptions between bouts of running.

We selected locomotion trials for which the average speed until the next locomotion offset point exceeded 1 cm/s and which lasted longer than 2 s. Quiescence trials were selected that lasted longer than 5 s, had an average speed <1 cm/s, and for which the maximum range of movement was <3 cm across the complete quiescence trial.

### Computation of LFP power

To compute LFP power spectra, we divided the data in 500 ms periods and multiplied each data segment by a Hann taper. We then computed the average LFP power spectrum by computing the FFT per segment and averaging over the segment’s power spectra.

### Spike-field locking analysis

Spike-field locking was computed using the Pairwise Phase Consistency (Vinck et al., 2012), a measure of phase consistency that is not biased by the firing rate or the number of spikes. The PPC is computed as follows:

1. For each spike, a spike-LFP phase is computed at each frequency (see below).
2. For each pair of spikes (fired by one cell) that fell in a different trial, we then compute the inner product of the two spike-LFP phases (the inner product being a measure of their similarity). Spike-LFP phases were computed for each spike and frequency separately by using Discrete Fourier Transform with Hanning taper of an LFP segment of length 9/*f*, where *f* is the frequency of interest. For a given time period (quiescence or locomotion), we only selected cells that fired at least 50 spikes in that period.
3. The PPC then equals the average of the inner products across all pairs of spike-LFP phases that fell in different trials. Note that exclusively taking pairs of spike-LFP phases from different trials ensures that history effects like bursting do not artificially inflate the measure of phase locking. The expected value of the PPC ranges between 0 and 1, although estimates lower than zero can occur.

For each cell, we computed the preferred phase of firing in the 40-60 Hz range by first computing the circular mean across all spike-LFP phases, and then computing the circular mean across frequencies. We then computed the phase consistency of preferred spike-LFP phases across units by computing the PPC over the preferred spike-LFP phases. Finally, we computed an estimate of the standard error of the mean using the jack-knife.

### Computation of STAs

To compute the average STA in the delta/theta [1,6] Hz and the gamma [40-60] Hz band, we first filtered the LFP data of 2s traces around each spike. We then normalized the energy of the LFP trace by either dividing by the mean absolute value of the LFP signal. We then averaged these traces across spikes.

### Computation of pair correlations

Unit-unit correlations were computed using the cross-correlogram at 1ms resolution. The cross-correlogram contains, for each cell pair, the number of spike coincidences at a certain delay (e.g. the number of times one neuron fired a spike 5-6 ms after another neuron). We normalized the cross-correlogram by computing the percent-wise increase compared to the expected fraction of coincidences given the firing rates of the two cells.

### Computation of modulation by state

To examine whether RS and FS cell firing rates were significantly changed around locomotion onset, we computed the firing rate in the [-0.5, 0.5] s window around locomotion onset (L-on; as in (Vinck et al., 2015) and compared this to the firing rate in the [-5,-2] s quiescence period before locomotion onset by computing log(FR_L-on_ / FR_Q_).

Using the same modulation index, we also compared the firing rate in the early [2,5] s quiescence period after locomotion offset – when the animal was still aroused but not yet moving (Vinck et al., 2015) – to the late [>40] s quiescence period after locomotion onset, when the animal had low arousal levels and is not moving (Vinck et al., 2015).

### Quantification of rate modulation to visual stimulus

For this analysis we computed the firing rate in the 30-500 ms period after stimulus onset, and the firing rate in the 1000 ms before stimulus onset. We then computed the firing rate modulation of stimulus-driven versus baseline rate as log(FR_stim_ / FR_base_). We call this modulation the SNR. We also computed the SNR at each time point by computing log(FR_stim_(t) / FR_base_) where FR_stim_(t) is the estimated firing rate at time *t*, which was estimated by convolving the spike trains with Gaussian smoothing windows (50ms, sigma=12.5ms), We further compared this modulation between the entire locomotion period and the entire quiescence period, computing the ΔSNR as log(FR_stim_, L / FR_base_, L) −log(FR_stim_, Q / FR_base_, Q).

### Quantification of visual response amplitude

For this analysis, we averaged the change in calcium fluorescence (ΔF/F(t)) in response to the three smallest contrasts (0-20%, C_low_), and in response to the three largest contrasts (90-100%, C_high_). We computed the contrast modulation as (C_high_-C_low_)/ (C_high_ + C_low_), and call this modulation the SNR. We determined the SNR during periods of quiescence and locomotion separately.

### Quantification of orientation selectivity

The orientation selectivity index is defined as R = [1-Circular Variance]. This is derived by letting each orientation (measured in radians, with 0 and 90 degree orientations corresponding to

0 and PI radians) be a vector on the circle with weight 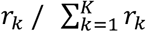. Note that we performed this procedure for the directions lying between 0 and 180 degrees and 180 and 360 degrees separately, and averaged all derived measures over the two set of directions. We then computed the resultant vector by summing the sine and cosine components.

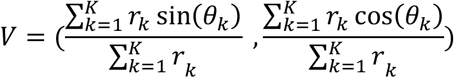

We next computed the resultant vector length (OSI) as R = |V|. Note that 0<=R<=1, and that R=1 indicates that a cell only has a non-zero firing rate for one orientation, whereas R=0 indicates that the cell has the same firing rates for all orientations. These vectors can also be normalized to unity length by V/|V|. These vectors are shown as a unit, and in histogram form. We also computed the phase consistency over the preferred orientations.

### Statistical comparisons

All statistics comparisons between groups were made using a permutation analysis. We first took the average value per animal and calculated group averages (control and mutant). For each permutation, condition labels (e.g., control versus mutant) were randomly permuted (shuffled) and the permuted data were used to generate group averages (control’ and mutant’). The difference in means between control’ and mutant’ was then calculated. This process was repeated 30,000 times to generate a null distribution for the group average difference in means. The p value for the actual difference in group means was calculated by summing the proportion of points beyond the actual value in the null distribution. For analyses requiring multiple comparisons across groups, we used a Bonferroni correction to adjust the threshold p value for significance. All permutation results are provided in Supplemental Table 1.

---

**Figure 1 Supplement.**
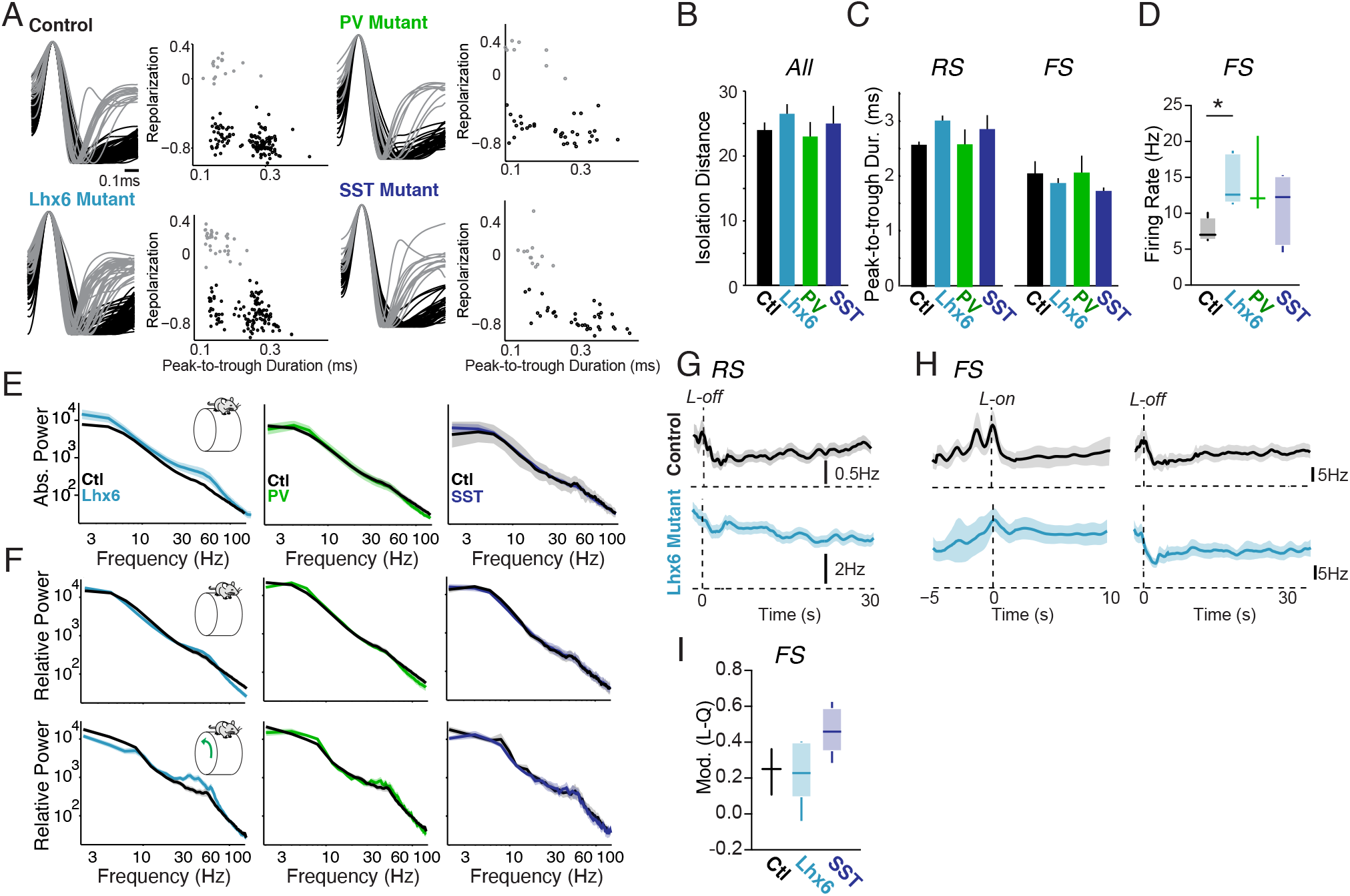
Waveform separation, firing rate, and LFP characteristics for ErbB4 mutants and controls. A. Normalized AP waveforms for controls and Lhx6, PV, and SST mutants, aligned to AP peak. Scatter plots show peak-to-trough duration of action potentials (APs) vs. AP repolarization metric for each group, which was defined as the value of the normalized (between −1 and +1) AP waveform at 450 ms (as in Vinck et al., 2015). Putative RS cells (black) can be subdivided into groups with thin and broad spikes (Haider et al., 2010; Vinck et al., 2015; Batista-Brito et al., 2017) that both have low firing rates in wild type animals (Vinck et al., 2015, Batista-Brito et al., 2017). Putative FS cells (gray) exhibit short duration and positive repolarization. B. Median ± s.e.m. of isolation distance for cells in mutant and control mice. C. Peak-to-trough duration for RS and FS cells. Error bars indicate mean ± s.e.m. D. Average firing rates for fast-spiking (FS) putative interneurons in controls and Lhx6 (cyan), PV (green), and SST (blue) mutants. Controls: 14 cells, 4 mice. Lhx6 mutants: 31 cells, 6 mice. PV mutants: 6 cells, 3 mice. SST mutants: 10 cells, 5 mice. E. Average firing rate for RS cells around locomotion offset (L-off) in controls (upper panel) and Lhx6 mutants (lower panel). F. Average firing rate for FS cells around locomotion onset (L-on; left) and offset (L-off; right) in controls (upper panel) and Lhx6 mutants (lower panel). G. Firing rate modulation index (L-Q/L+Q) in early locomotion period (L; −0.5 to 0.5s around L-on) as compared to quiescence period (Q) for FS cells in each group. Note that the number of FS cell recordings during state transitions in the PV mutant mice was insufficient for statistical comparisons, and these data were therefore not included here. Controls: 11 cells, 3 mice. Lhx6 mutants: 27 cells, 6 mice. SST mutants: 9 cells, 5 mice. H. LFP power spectra during quiescence in controls and Lhx6, PV, and SST mutants. I. LFP power spectra, shown as relative power, for quiescence (upper panels) and locomotion (lower panels) in each group.

**Figure 2 Supplement.**
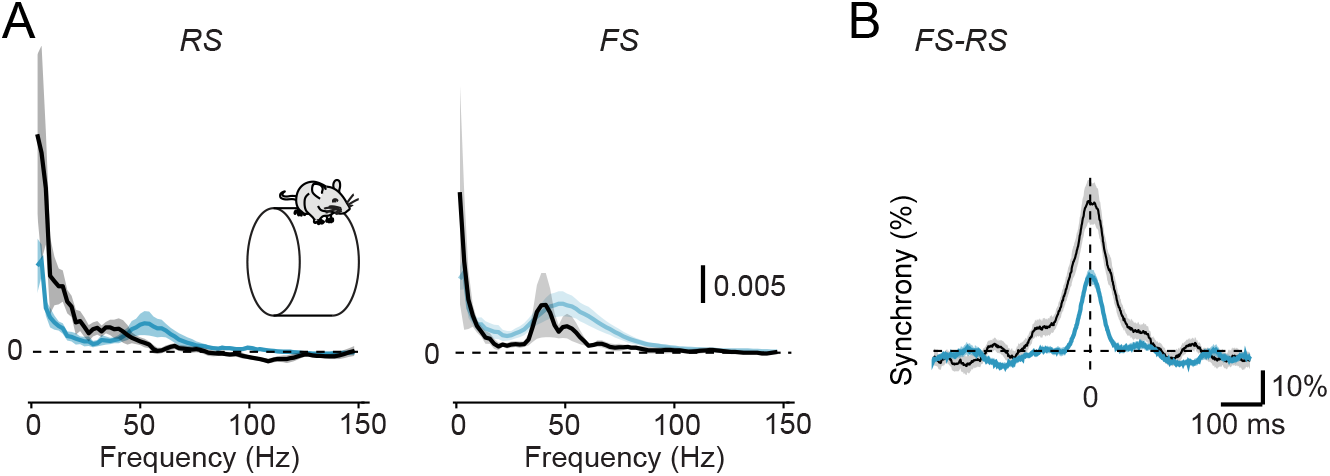
Disrupted temporal organization of cortical activity. A. Average spike-LFP phase locking during quiescent periods for RS (left) and FS (right) cells in controls (black) and Lhx6 mutants (cyan). B. Average normalized cross-correlogram during quiescence for RS-FS pairs. (Controls: 79 RS-FS pairs, 5 mice. Lhx6 mutants: 6 mice)

**Figure 3 Supplement.**
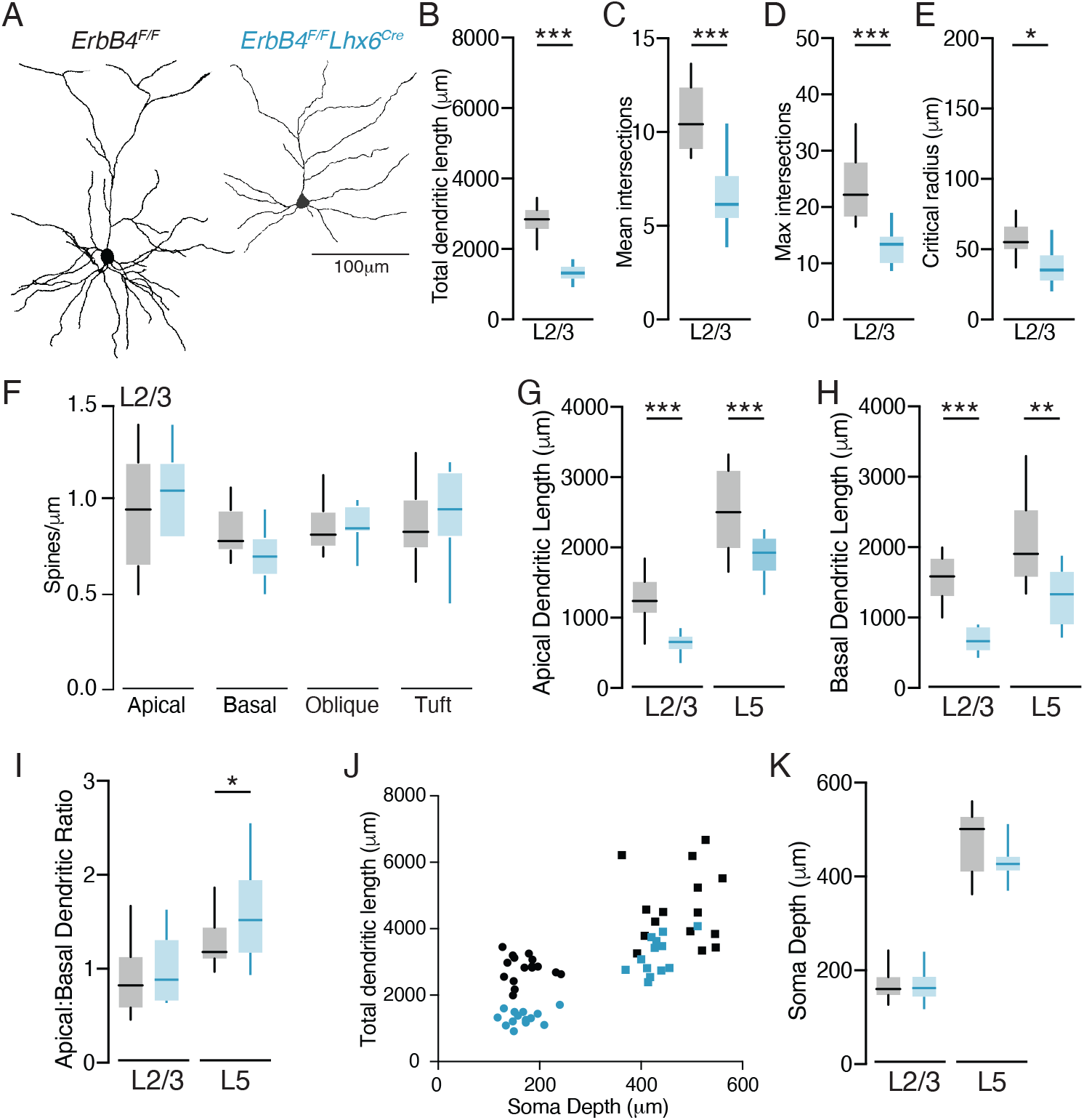
Morphological characteristics of pyramidal neurons in ErbB4 mutants. A. Example layer 3 pyramidal neuron reconstructions from an *ErbB4^F/F^* control (left) and an *ErbB4^F/F^, Lhx6^Cre^* mutant (right). B. Average total dendritic length for layer 3 pyramidal neurons in controls (black) and Lhx6 mutants (cyan). Controls: 16 cells. Lhx6 mutants: 14 cells. C. Average mean dendritic intersections for control and Lhx6 mutant layer 3 cells. D. Average maximum dendritic intersections for control and Lhx6 mutant layer 3 cells. E. Average critical radius for control and Lhx6 mutant layer 3 cells. F. Spine density on the main apical dendrite, basal dendrites, oblique dendrites, and apical dendritic tuft for layer 3 pyramidal neurons in controls and Lhx6 mutants. Controls: 10 cells. Lhx6 mutants: 10 cells. G. Scatter plot of soma depth vs dendritic length for all pyramidal neurons reconstructed in controls and Lhx6 mutants. H. Average apical dendritic length for layer 2/3 and layer 5 neurons in controls and Lhx6 mutants. I. Average basal dendritic length for layer 2/3 and layer 5 neurons in controls and Lhx6 mutants. Average apical:basal dendritic length ratio for layer 2/3 and layer 5 neurons in controls and Lhx6 mutants. K. Average soma depth of all reconstructed neurons in controls and mutants.

**Figure 4 Supplement.**
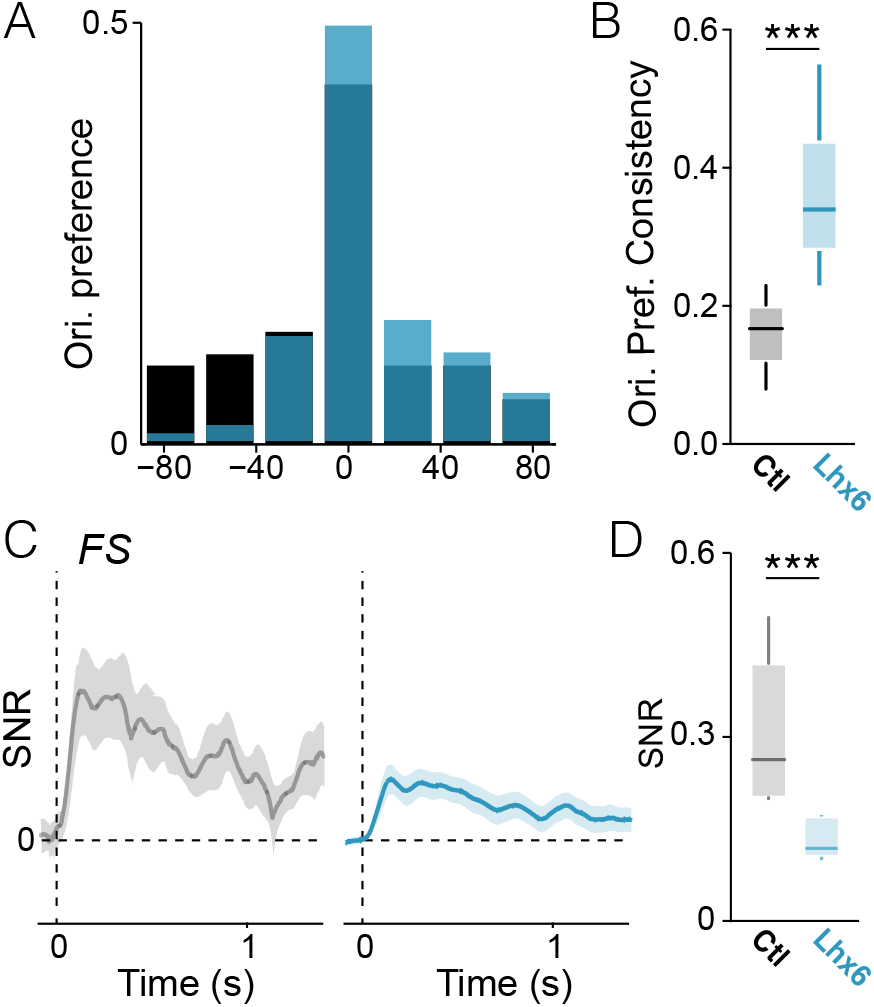
Additional quantification of visual responses in ErbB4 mutants. A. Histogram of preferred orientations across the population of RS cells in controls (black) and Lhx6 mutants (cyan). B. Population phase consistency of orientation preferences for controls and mutants. Both controls and mutants exhibit significant clustering of orientation preferences, but mutants show more clustering than controls. C. Average visual response of FS cells in each group, shown as signal:noise ratio over the stimulus presentation period (SNR). D. Average signal-to-noise ratio of visual responses for FS cells in controls and Lhx6 mutants. Controls: 11 cells, 5 mice. Lhx6 mutants: 22 cells, 5 mice.

**Supplemental Table 1.**
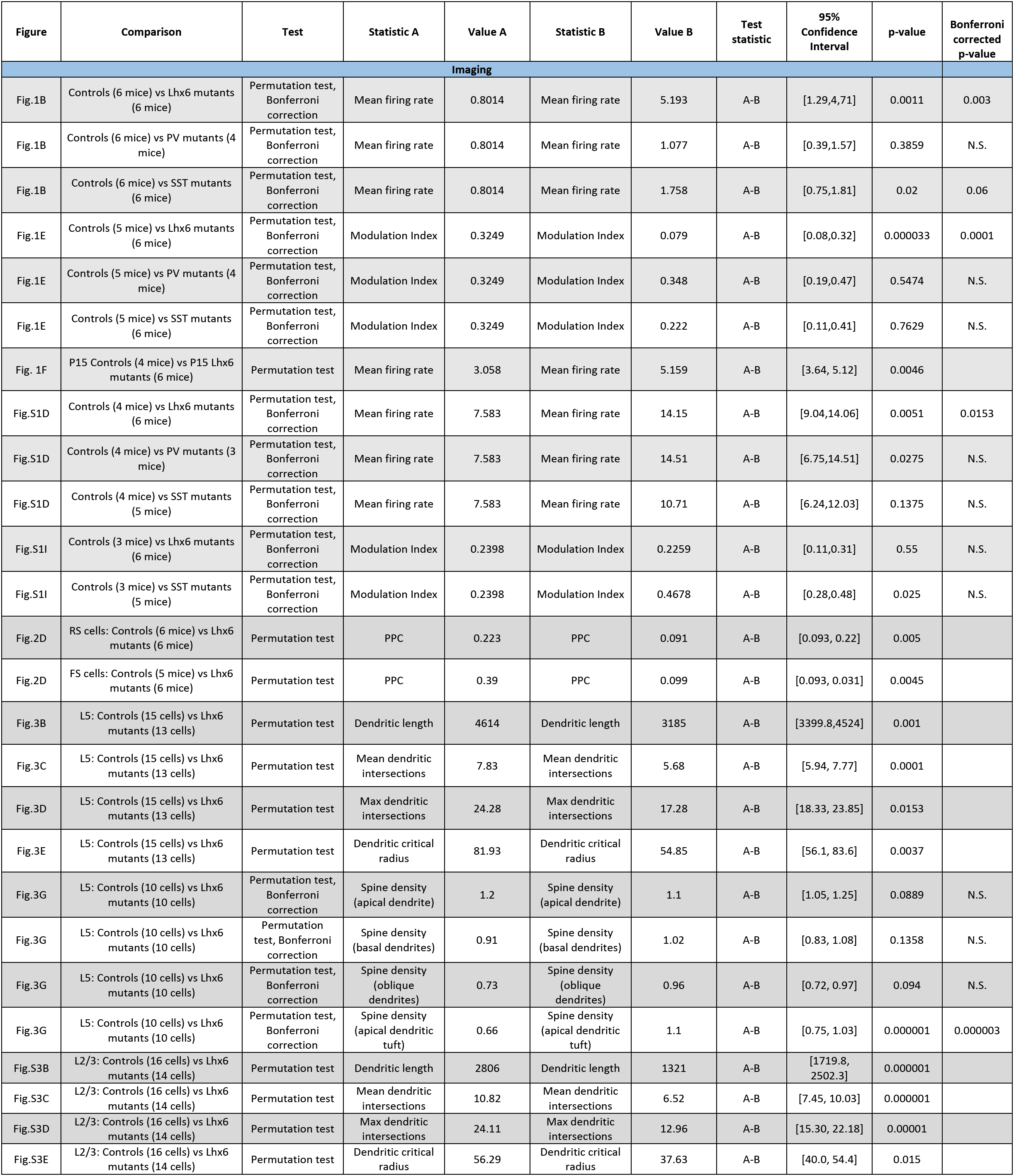
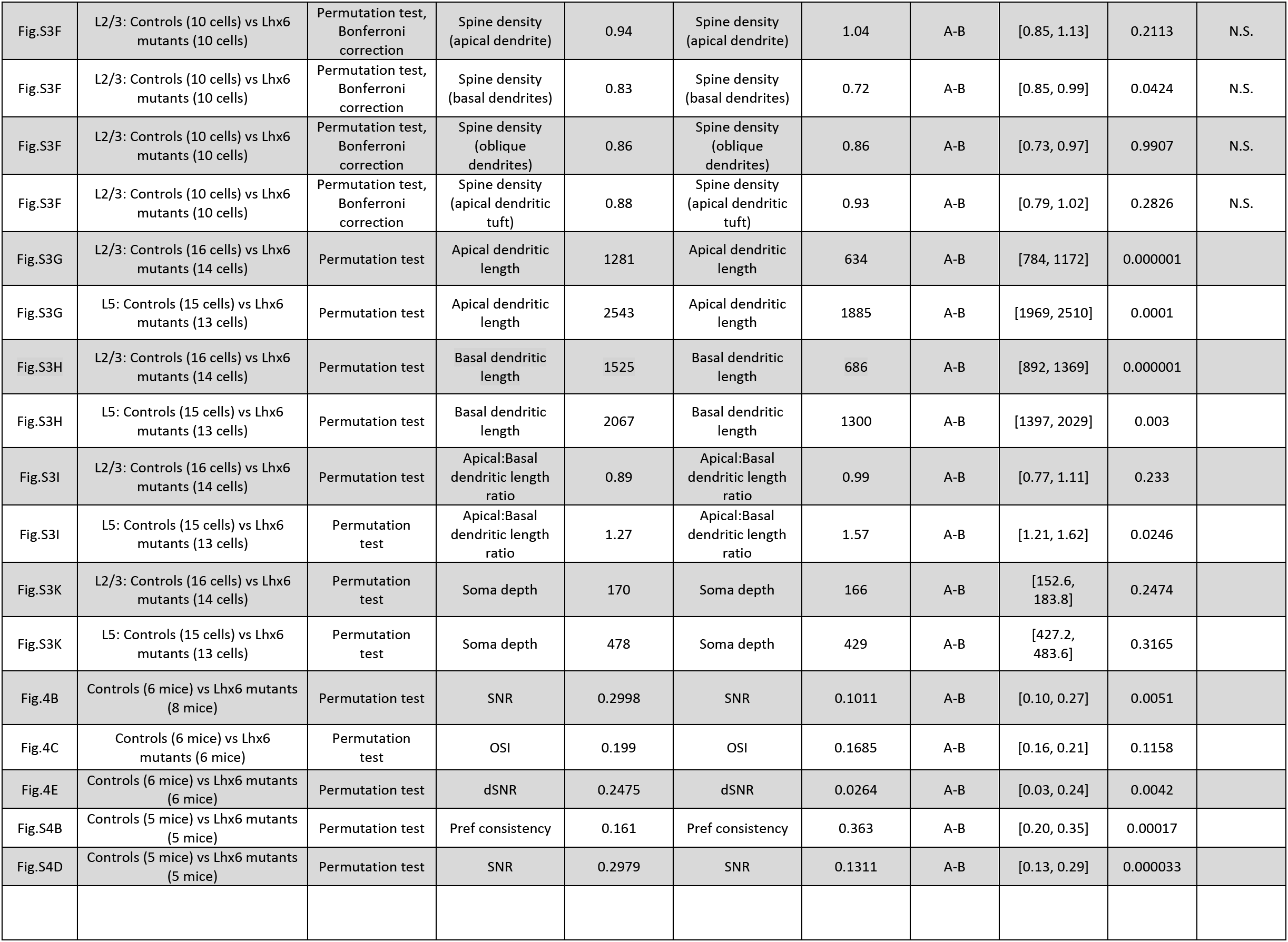
Summary of all statistical analyses.

## Notes

### Competing Interest Statement

The authors have declared no competing interest.

## References

Abe Y, Namba H, Kato T, Iwakura Y, Nawa H (2011) Neuregulin-1 signals from the periphery regulate AMPA receptor sensitivity and expression in GABAergic interneurons in developing neocortex. J Neurosci 31:5699–5709.

Allene C, Cattani A, Ackman JB, Bonifazi P, Aniksztejn L, Ben-Ari Y, Cossart R (2008) Sequential generation of two distinct synapse-driven network patterns in developing neocortex. J Neurosci 28:12851–12863.

Barros CS, Calabrese B, Chamero P, Roberts AJ, Korzus E, Lloyd K, Stowers L, Mayford M, Halpain S, Muller U (2009) Impaired maturation of dendritic spines without disorganization of cortical cell layers in mice lacking NRG1/ErbB signaling in the central nervous system. Proc Natl Acad Sci USA 106:4507–4512.

Batista-Brito R, Vinck M, Ferguson KA, Chang JT, Laubender D, Lur G, Mossner JM, Hernandez VG, Ramakrishnan C, Deisseroth K, Higley MJ, Cardin JA (2017) Developmental Dysfunction of VIP Interneurons Impairs Cortical Circuits. Neuron 95:884–895 e889.

Bennett C, Arroyo S, Hestrin S (2013) Subthreshold mechanisms underlying state-dependent modulation of visual responses. Neuron 80:350–357.

Cardin JA (2016) Snapshots of the Brain in Action: Local Circuit Operations through the Lens of gamma Oscillations. The Journal of neuroscience : the official journal of the Society for Neuroscience 36:10496–10504.

Cardin JA (2018) Inhibitory Interneurons Regulate Temporal Precision and Correlations in Cortical Circuits. Trends in neurosciences 41:689–700.

Cardin JA, Kumbhani RD, Contreras D, Palmer LA (2010) Cellular mechanisms of temporal sensitivity in visual cortex neurons. The Journal of neuroscience : the official journal of the Society for Neuroscience 30:3652–3662.

Cardin JA, Carlen M, Meletis K, Knoblich U, Zhang F, Deisseroth K, Tsai LH, Moore CI (2009) Driving fast-spiking cells induces gamma rhythm and controls sensory responses. Nature 459:663–667.

Cauller LJ, Clancy B, Connors BW (1998) Backward cortical projections to primary somatosensory cortex in rats extend long horizontal axons in layer I. J Comp Neurol 390:297–310.

Chen YJ, Zhang M, Yin DM, Wen L, Ting A, Wang P, Lu YS, Zhu XH, Li SJ, Wu CY, Wang XM, Lai C, Xiong WC, Mei L, Gao TM (2010) ErbB4 in parvalbumin-positive interneurons is critical for neuregulin 1 regulation of long-term potentiation. Proc Natl Acad Sci U S A 107:21818–21823.

Chiu CQ, Lur G, Morse TM, Carnevale NT, Ellis-Davies GC, Higley MJ (2013) Compartmentalization of GABAergic inhibition by dendritic spines. Science 340:759–762.

Cossart R (2011) The maturation of cortical interneuron diversity: how multiple developmental journeys shape the emergence of proper network function. Curr Opin Neurobiol 21:160–168.

Cruz-Martin A, El-Danaf RN, Osakada F, Sriram B, Dhande OS, Nguyen PL, Callaway EM, Ghosh A, Huberman AD (2014) A dedicated circuit links direction-selective retinal ganglion cells to the primary visual cortex. Nature 507:358–361.

David C, Schleicher A, Zuschratter W, Staiger JF (2007) The innervation of parvalbumin-containing interneurons by VIP-immunopositive interneurons in the primary somatosensory cortex of the adult rat. Eur J Neurosci 25:2329–2340.

Del Pino I, Brotons-Mas JR, Marques-Smith A, Marighetto A, Frick A, Marin O, Rico B (2017) Abnormal wiring of CCK(+) basket cells disrupts spatial information coding. Nat Neurosci 20:784–792.

Del Pino I, Garcia-Frigola C, Dehorter N, Brotons-Mas JR, Alvarez-Salvado E, Martinez de Lagran M, Ciceri G, Gabaldon MV, Moratal D, Dierssen M, Canals S, Marin O, Rico B (2013) Erbb4 Deletion from Fast-Spiking Interneurons Causes Schizophrenia-like Phenotypes. Neuron 79:1152–1168.

del Rio JA, de Lecea L, Ferrer I, Soriano E (1994) The development of parvalbumin-immunoreactivity in the neocortex of the mouse. Brain research Developmental brain research 81:247–259.

Dipoppa M, Ranson A, Krumin M, Pachitariu M, Carandini M, Harris KD (2018) Vision and Locomotion Shape the Interactions between Neuron Types in Mouse Visual Cortex. Neuron 98:602–615 e608.

Fazzari P, Paternain AV, Valiente M, Pla R, Lujan R, Lloyd K, Lerma J, Marin O, Rico B (2010) Control of cortical GABA circuitry development by Nrg1 and ErbB4 signalling. Nature 464:1376–1380.

Felleman DJ, Van Essen DC (1991) Distributed hierarchical processing in the primate cerebral cortex. Cereb Cortex 1:1–47.

Fishell G, Rudy B (2011) Mechanisms of inhibition within the telencephalon: “where the wild things are”. Annu Rev Neurosci 34:535–567.

Fogarty M, Grist M, Gelman D, Marin O, Pachnis V, Kessaris N (2007) Spatial genetic patterning of the embryonic neuroepithelium generates GABAergic interneuron diversity in the adult cortex. J Neurosci 27:10935–10946.

Fries P (2009) Neuronal gamma-band synchronization as a fundamental process in cortical computation. Annual review of neuroscience 32:209–224.

Fu Y, Tucciarone JM, Espinosa JS, Sheng N, Darcy DP, Nicoll RA, Huang ZJ, Stryker MP (2014a) A cortical circuit for gain control by behavioral state. Cell 156:1139–1152.

Fu Y, Tucciarone JM, Espinosa JS, Sheng N, Darcy DP, Nicoll RA, Huang ZJ, Stryker MP (2014b) A cortical circuit for gain control by behavioral state. Cell 156:1139–1152.

Gandal MJ, Edgar JC, Klook K, Siegel SJ (2012) Gamma synchrony: towards a translational biomarker for the treatment-resistant symptoms of schizophrenia. Neuropharmacology 62:1504–1518.

Gibson JR, Beierlein M, Connors BW (1999) Two networks of electrically coupled inhibitory neurons in neocortex. Nature 402:75–79.

Goldberg EM, Jeong HY, Kruglikov I, Tremblay R, Lazarenko RM, Rudy B (2011) Rapid developmental maturation of neocortical FS cell intrinsic excitability. Cereb Cortex 21:666–682.

Haider B, Krause MR, Duque A, Yu Y, Touryan J, Mazer JA, McCormick DA (2010) Synaptic and network mechanisms of sparse and reliable visual cortical activity during nonclassical receptive field stimulation. Neuron 65:107–121.

Hasenstaub A, Shu Y, Haider B, Kraushaar U, Duque A, McCormick DA (2005) Inhibitory postsynaptic potentials carry synchronized frequency information in active cortical networks. Neuron 47:423–435.

Jia H, Rochefort NL, Chen X, Konnerth A (2010) Dendritic organization of sensory input to cortical neurons in vivo. Nature 464:1307–1312.

Keller AJ, Roth MM, Scanziani M (2020) Feedback generates a second receptive field in neurons of the visual cortex. Nature 582:545–549.

Ko H, Hofer SB, Pichler B, Buchanan KA, Sjostrom PJ, Mrsic-Flogel TD (2011) Functional specificity of local synaptic connections in neocortical networks. Nature 473:87–91.

Larkum ME, Zhu JJ (2002) Signaling of layer 1 and whisker-evoked Ca2+ and Na+ action potentials in distal and terminal dendrites of rat neocortical pyramidal neurons in vitro and in vivo. J Neurosci 22:6991–7005.

Larkum ME, Senn W, Luscher HR (2004) Top-down dendritic input increases the gain of layer 5 pyramidal neurons. Cereb Cortex 14:1059–1070.

Lavdas AA, Grigoriou M, Pachnis V, Parnavelas JG (1999) The medial ganglionic eminence gives rise to a population of early neurons in the developing cerebral cortex. J Neurosci 19:7881–7888.

Lewis DA, Lieberman JA (2000) Catching up on schizophrenia: natural history and neurobiology. Neuron 28:325–334.

Lien AD, Scanziani M (2013) Tuned thalamic excitation is amplified by visual cortical circuits. Nat Neurosci 16:1315–1323.

McGinley MJ, Vinck M, Reimer J, Batista-Brito R, Zagha E, Cadwell CR, Tolias AS, Cardin JA, McCormick DA (2015) Waking State: Rapid Variations Modulate Neural and Behavioral Responses. Neuron 87:1143–1161.

Modol L, Sousa VH, Malvache A, Tressard T, Baude A, Cossart R (2017) Spatial Embryonic Origin Delineates GABAergic Hub Neurons Driving Network Dynamics in the Developing Entorhinal Cortex. Cereb Cortex 27:4649–4661.

Mossner JM, Batista-Brito R, Pant R, Cardin JA (2020) Developmental loss of MeCP2 from VIP interneurons impairs cortical function and behavior. eLife 9.

Nassi JJ, Lomber SG, Born RT (2013) Corticocortical feedback contributes to surround suppression in V1 of the alert primate. J Neurosci 33:8504–8517.

Neddens J, Buonanno A (2010) Selective populations of hippocampal interneurons express ErbB4 and their number and distribution is altered in ErbB4 knockout mice. Hippocampus 20:724–744.

Niell CM, Stryker MP (2010) Modulation of visual responses by behavioral state in mouse visual cortex. Neuron 65:472–479.

Pakan JM, Lowe SC, Dylda E, Keemink SW, Currie SP, Coutts CA, Rochefort NL (2016) Behavioral-state modulation of inhibition is context-dependent and cell type specific in mouse visual cortex. eLife 5.

Pfeffer CK, Xue M, He M, Huang ZJ, Scanziani M (2013) Inhibition of inhibition in visual cortex: the logic of connections between molecularly distinct interneurons. Nat Neurosci 16:1068–1076.

Pi HJ, Hangya B, Kvitsiani D, Sanders JI, Huang ZJ, Kepecs A (2013) Cortical interneurons that specialize in disinhibitory control. Nature 503:521–524.

Pouille F, Scanziani M (2001) Enforcement of temporal fidelity in pyramidal cells by somatic feed-forward inhibition. Science 293:1159–1163.

Ranganathan GN, Apostolides PF, Harnett MT, Xu NL, Druckmann S, Magee JC (2018) Active dendritic integration and mixed neocortical network representations during an adaptive sensing behavior. Nat Neurosci 21:1583–1590.

Rochefort NL, Narushima M, Grienberger C, Marandi N, Hill DN, Konnerth A (2011) Development of direction selectivity in mouse cortical neurons. Neuron 71:425–432.

Rockland KS, Pandya DN (1979) Laminar origins and terminations of cortical connections of the occipital lobe in the rhesus monkey. Brain Res 179:3–20.

Rossignol E (2011) Genetics and function of neocortical GABAergic interneurons in neurodevelopmental disorders. Neural plasticity 2011:649325.

Roth MM, Dahmen JC, Muir DR, Imhof F, Martini FJ, Hofer SB (2016) Thalamic nuclei convey diverse contextual information to layer 1 of visual cortex. Nat Neurosci 19:299–307.

Schiller J, Schiller Y, Stuart G, Sakmann B (1997) Calcium action potentials restricted to distal apical dendrites of rat neocortical pyramidal neurons. J Physiol 505 (Pt 3):605–616.

Takesian AE, Hensch TK (2013) Balancing plasticity/stability across brain development. Progress in brain research 207:3–34.

Takesian AE, Bogart LJ, Lichtman JW, Hensch TK (2018) Inhibitory circuit gating of auditory critical-period plasticity. Nat Neurosci 21:218–227.

Tan Z, Robinson HL, Yin DM, Liu Y, Liu F, Wang H, Lin TW, Xing G, Gan L, Xiong WC, Mei L (2018) Dynamic ErbB4 Activity in Hippocampal-Prefrontal Synchrony and Top-Down Attention in Rodents. Neuron 98:380–393 e384.

Tang L, Higley MJ (2020) Layer 5 Circuits in V1 Differentially Control Visuomotor Behavior. Neuron 105:346–354 e345.

Ting AK, Chen Y, Wen L, Yin DM, Shen C, Tao Y, Liu X, Xiong WC, Mei L (2011) Neuregulin 1 promotes excitatory synapse development and function in GABAergic interneurons. J Neurosci 31:15–25.

Tuncdemir SN, Wamsley B, Stam FJ, Osakada F, Goulding M, Callaway EM, Rudy B, Fishell G (2016) Early Somatostatin Interneuron Connectivity Mediates the Maturation of Deep Layer Cortical Circuits. Neuron 89:521–535.

Uhlhaas PJ, Singer W (2010) Abnormal neural oscillations and synchrony in schizophrenia. Nature reviews Neuroscience 11:100–113.

Urban-Ciecko J, Jouhanneau JS, Myal SE, Poulet JFA, Barth AL (2018) Precisely Timed Nicotinic Activation Drives SST Inhibition in Neocortical Circuits. Neuron 97:611–625 e615.

Vinck M, Battaglia FP, Womelsdorf T, Pennartz C (2012) Improved measures of phase-coupling between spikes and the Local Field Potential. Journal of computational neuroscience 33:53–75.

Vinck M, Batista-Brito R, Knoblich U, Cardin JA (2015) Arousal and locomotion make distinct contributions to cortical activity patterns and visual encoding. Neuron 86:740–754.

Vinck M, Womelsdorf T, Buffalo EA, Desimone R, Fries P (2013) Attentional modulation of cell-class-specific gamma-band synchronization in awake monkey area v4. Neuron 80:1077–1089.

Vullhorst D, Neddens J, Karavanova I, Tricoire L, Petralia RS, McBain CJ, Buonanno A (2009) Selective expression of ErbB4 in interneurons, but not pyramidal cells, of the rodent hippocampus. J Neurosci 29:12255–12264.

Walker F, Mock M, Feyerabend M, Guy J, Wagener RJ, Schubert D, Staiger JF, Witte M (2016) Parvalbumin- and vasoactive intestinal polypeptide-expressing neocortical interneurons impose differential inhibition on Martinotti cells. Nature communications 7:13664.

Wang C, Waleszczyk WJ, Burke W, Dreher B (2007) Feedback signals from cat’s area 21a enhance orientation selectivity of area 17 neurons. Exp Brain Res 182:479–490.

Wehr M, Zador AM (2003) Balanced inhibition underlies tuning and sharpens spike timing in auditory cortex. Nature 426:442–446.

Yau HJ, Wang HF, Lai C, Liu FC (2003) Neural development of the neuregulin receptor ErbB4 in the cerebral cortex and the hippocampus: preferential expression by interneurons tangentially migrating from the ganglionic eminences. Cereb Cortex 13:252–264.

Yin DM, Chen YJ, Lu YS, Bean JC, Sathyamurthy A, Shen C, Liu X, Lin TW, Smith CA, Xiong WC, Mei L (2013) Reversal of behavioral deficits and synaptic dysfunction in mice overexpressing neuregulin 1. Neuron 78:644–657.

Zhang S, Xu M, Kamigaki T, Hoang Do JP, Chang WC, Jenvay S, Miyamichi K, Luo L, Dan Y (2014) Selective attention. Long-range and local circuits for top-down modulation of visual cortex processing. Science 345:660–665.

